# Independent Mesh Realizations Introduce Percent-Level Variability in Temporal Interference Simulations

**DOI:** 10.64898/2026.08.08.743658

**Authors:** Boyan Ivanov, Mahnaz Arvaneh, Jake Toth, Sumientra Rampersad

## Abstract

Computational models of temporal interference stimulation (TIS) commonly report a single electric-field estimate for a given anatomy and electrode montage. Because non-deterministic tetrahedral mesh generation does not produce a unique discretisation of a fixed tissue-label image, a single mesh realisation may introduce numerical variability. We quantified variation across independent mesh realisations and contrasted it with repeated downstream simulation execution on a single selected mesh. Ten head models were evaluated for stimulation of the left hippocampus and right primary motor cortex (M1). For every model and target, we generated 40 independent meshes and performed one complete simulation on each. Separately, we selected the mesh whose parcel-level field estimate was closest to the median and repeated downstream operations 40 times while holding that geometry fixed, yielding 1,600 TIS simulations in total. The primary outcome was the spatial median of the TIS envelope field within a spherical target region. Across independently remeshed runs, within-participant coefficients of variation were 1.81–3.65% for the hippocampus and 1.62–2.79% for M1. Repeated execution on a fixed mesh reduced run-to-run standard deviation by more than 99%, demonstrating that workflow variability is driven almost entirely by non-deterministic mesh generation rather than solver instability, numerical rounding, or post-processing. Single-run mesh realisations preserved overall cohort ordering (median Kendall’s *τ* of 0.867 for the hippocampus and 0.911 for M1) but frequently inverted the rank order of participant pairs with similar predicted fields. Furthermore, a bootstrap analysis demonstrated that averaging five to ten independent remesh runs effectively suppressed this stochastic noise. These results quantify single-workflow repeatability rather than absolute error. Stochastic mesh variation should therefore be controlled or mitigated through multi-run averaging whenever experimental conclusions depend on subtle field differences or fixed neuromodulation thresholds.

## 1 Introduction

Transcranial temporal interference stimulation (TIS) is a non-invasive brain stimulation method that applies two high-frequency electric fields with a small frequency difference, producing a lower-frequency modulation envelope where the fields overlap [1]. Experiments in animals and humans have demonstrated that TIS can modulate neural activity and produce measurable behavioural effects [1, 2, 3]. Computational modelling is central to the design and interpretation of TIS. Models estimate the spatial distribution of the TIS envelope field and support the selection of optimal electrode montages for a chosen brain target [1, 4]. The finite element method (FEM) is commonly used because it can represent the irregular geometry and heterogeneous conductivities of the head. FEM divides the volume conductor into a mesh of tetrahedral elements, applies appropriate boundary conditions, and then approximates the electric potential over this discrete geometry [5, 6]. The resulting field therefore depends on both the physical head model and its numerical representation. Head-mesh generation introduces a source of numerical uncertainty that is distinct from segmentation or conductivity errors. Standard finite-element modelling packages for non-invasive brain stimulation, such as SimNIBS [7] and ROAST [8], employ non-deterministic meshing algorithms. Consequently, a single mesh generation step yields one valid discretisation of a tissue-label image rather than a unique numerical ground truth. A new tetrahedralisation occurs whenever the tissue-label image or meshing parameters change, for example after correcting a segmentation or altering the mesh resolution. In this study, mesh generation was deliberately repeated between otherwise identical runs with unchanged settings to quantify variability among valid discretisations of the same underlying anatomy. Independent meshes can differ in their element boundaries, element counts, and discretised tissue volumes. These differences alter the numerical approximation of the volume-conduction problem. Consequently, a change in the predicted field between simulations does not necessarily result from a change in anatomy or assigned conductivity values.

Mesh-related uncertainty also matters when only one simulation is performed for a given tissue-label image. In that common situation, the reported result represents an arbitrary point sample from an underlying distribution of valid discretisations. Small differences in calculated fields between electrode configurations or participants may therefore reflect stochastic mesh realisation rather than true anatomical or physiological differences. This stochastic variation can have important practical consequences when interpreting simulation results. First, when participants are ranked by predicted target field strength to compare individual responses or customise dosing, meshing noise between closely matched individuals can artificially reverse their relative rank order. Second, when researchers evaluate target exposure using a defined electric field threshold (for example, determining whether a region receives at least 0.20 V/m of electric field to infer potential neural engagement [4, 9]), numerical variation between independent mesh realisations can incorrectly push a participant’s estimate above or below that cutoff value.

Previous work in transcranial electric field modelling has extensively evaluated sources of uncertainty such as segmentation accuracy, tissue conductivity variability, electrode placement, and finite-element solver convergence [5, 10, 4, 6, 7]. While mesh-convergence studies test whether field estimates stabilise as element resolution increases, they assume that a given meshing algorithm produces deterministic outputs at a specified density. Consequently, the baseline numerical uncertainty introduced purely by non-deterministic, stochastic remeshing of an identical tissue-label image remains unquantified, both for temporal interference stimulation and for transcranial electric modelling more broadly.

Although stochastic tetrahedralisation is an inherent feature of finite-element modelling pipelines across electrical stimulation modalities, TIS provides a unique paradigm to evaluate this phenomenon. First, unlike other modalities, TIS requires solving two independent electric field problems on the base mesh and combining them into a spatial modulation envelope that depends on local field magnitudes, vector orientations, and piecewise conditional bounds [1, 4]. Second, while conventional transcranial stimulation is primarily applied to superficial sites, human experimental TIS protocols are actively deployed to target both superficial cortical regions, such as the primary motor cortex [11, 12], and deep subcortical structures, such as the hippocampus or striatum [2, 3]. Evaluating meshing variability across both superficial and deep target geometries therefore directly addresses the dual clinical landscape in which TIS computational models are actually utilised.

In this study, we addressed two main research questions through TIS simulations targeting the left hippocampus or right primary motor cortex (M1) across ten head models. First, how much variability arises across independently generated base head meshes compared with repeated execution of downstream simulation steps on one selected fixed mesh? Second, how many independent mesh realizations must be averaged to stabilize this variability and achieve a reliable target-field estimate? We evaluated this using a bootstrap analysis to quantify how sampling precision improves as additional remeshed runs are included. Element-count and tissue-composition analyses were used to characterise how the underlying discretisations changed between independent runs.

## 2 Methods

### 2.1 Study design

This study completed 40 runs of the same workflow under two numerical conditions across 10 head models and two brain targets. A single workflow run comprised construction of the stimulation montage on a base head mesh, simulation of the two component electric fields, calculation of the temporal interference envelope field (hereafter referred to as the *TIS field*), interpolation of the field to voxel space, and target-field summarisation (where *target field* denotes the TIS field evaluated within a defined target region). The two conditions differed in how the base mesh was constructed. In the *remesh condition*, a new tetrahedral head mesh was generated from the same tissue-label image before each run. In the *fixed-mesh condition*, one mesh was selected from the corresponding remesh distribution and loaded for every run. The study design is summarised in Table 1. The comparison contrasts repeated-run variability in the complete workflow (which included remeshing) with downstream repeatability conditional on one selected fixed mesh. Because downstream repetitions were not performed across multiple randomly sampled meshes, the design does not provide a formal separation of between-mesh and within-mesh variance, does not identify which mesh is closest to the continuum solution, and does not yield a complete variance decomposition across multiple fixed meshes.

**Table 1:** Study structure. The study design separates mesh-generation variability from downstream simulation variability.

| Condition | Runs per subject | Mesh handling | Variation represented |
| --- | --- | --- | --- |
| Remesh | 40 | CHARM generated a new head mesh from the same corrected tissue labels before every run | Variations caused by mesh generation and all downstream simulation steps |
| Fixed mesh | 40 | All runs reused one fixed subject-specific head mesh | Variations caused only by the downstream simulation steps |

Total: 10 participants × 2 targets × 2 conditions × 40 runs = 1,600 TIS simulations.

Forty runs per condition were selected pragmatically to characteristically represent the run-to-run distribution across many realisations while keeping computational demand feasible. The resulting dataset supported a retrospective bootstrap analysis of averages containing 2 to 40 independently meshed runs to evaluate how many runs are required to achieve a stable target-field estimate.

For the remesh condition, the hippocampal and primary motor cortex (M1) experiments used independently generated collections of 40 meshes per participant. For the fixed-mesh condition, one mesh was selected for each participant–target combination post hoc following the 40 remesh runs. Mesh selection was based on the median TIS field across the complete anatomical target parcel. The run whose parcel-level value was closest to the median of the corresponding 40-run distribution supplied the saved mesh used in all fixed-mesh runs. This data-dependent selection rule used a larger region than the spherical primary ROI defined below and was intended to avoid choosing an extreme mesh realisation rather than to establish an independent reference geometry. Across 10 participants, two targets, two numerical conditions, and 40 runs per condition, the study comprised 1,600 TIS simulations.

### 2.2 Study population and imaging data

Ten datasets from the Cambridge Centre for Ageing and Neuroscience database formed the biological sample [13]. Participants were selected to balance recorded sex and span the adult age range. Five were recorded as female and five as male. Median age was 52.58 years, with a range of 26.00–79.17 years (Table 2).

**Table 2:** Study population. Demographic details of the 10 CamCAN participants used in this study. Participants were selected randomly such that it balances sex and age range across the cohort.

| CamCAN ID | Recorded sex | Age (years) |
| --- | --- | --- |
| CC110174 | Female | 26.00 |
| CC121144 | Male | 26.00 |
| CC310407 | Female | 39.00 |
| CC320616 | Male | 39.08 |
| CC420071 | Female | 52.58 |
| CC410432 | Male | 52.58 |
| CC520083 | Female | 65.08 |
| CC520127 | Male | 66.00 |
| CC610631 | Female | 77.58 |
| CC720941 | Male | 79.17 |

Each dataset comprised high-resolution structural MRI acquired using a 3 T Siemens MAGNETOM TrioTim scanner equipped with a 32-channel head coil. T1-weighted images were acquired using a sagittal three-dimensional magnetisation-prepared rapid gradient-echo sequence with a repetition time (TR) of 2250 ms, an echo time (TE) of 2.98 ms, an inversion time of 900 ms, a flip angle of 9 degrees, and a voxel size of 1 x 1 x 1 mm³. T2-weighted images were acquired using a sagittal three-dimensional variable-flip-angle SPACE sequence with a TR of 2800 ms, a TE of 408 ms, and a voxel size of 1 x 1 x 1 mm³.

### 2.3 Segmentation and meshing

Initial tissue segmentation labels were generated from the structural images using CHARM [6, 7]. The label map represented scalp, compact and spongy bone, cerebrospinal fluid (CSF), grey matter, white matter, eyes, muscle, and blood. Tissue-label corrections were performed automatically using connected-component filtering and binary morphology. The largest 26-connected component was retained first for white matter and then for the combined white- and grey-matter mask. Blood was excluded temporarily while the combined white-matter, grey-matter, and CSF envelope was corrected. The largest 26-connected component of this envelope was closed with a spherical structuring element of radius 7 voxels, opened with a radius of 3 voxels to remove narrow protrusions, and filtered once more to retain its largest component. Blood and the separate tissue labels were then restored. Gaps in the outer head boundary were filled by closing the complete tissue envelope with a spherical structuring element of radius 10 voxels. One additional binary-closing iteration with a 3 x 3 x 3-voxel structuring element was applied to the scalp label.

An initial test mesh was generated in CHARM for quality control, and all tissue surfaces were reviewed by an expert to ensure anatomical validity and rule out topological defects. Once a participant’s head model passed inspection, this initial test mesh was discarded, and the corrected voxelwise tissue-label map was saved. For each participant, this tissue-label image was used as the input for the mesh-generation step of every subsequent remesh run.

### 2.4 Regions of interest

Montage optimization and outcome evaluation used anatomically constrained spherical regions of interest (ROIs). The same ROI-construction procedure was applied to the MNI152 model used for optimization and to each of the 10 participant datasets. FreeSurfer 7.4.1 processing identified the volumetric left hippocampus and the right precentral-gyrus parcel on each T1-weighted image using the Destrieux atlas [14, 15]. The right-M1 target was operationalised using the right precentral-gyrus parcel. The volumetric centroid of each parcel defined the centre of a sphere, which was clipped to the boundaries of the corresponding anatomical parcel. The sphere radius was increased until the clipped region reached the specified target volume. Different target volumes were used for the hippocampus and M1 ROI to account for the thin, sheet-like geometry of the cortex. The resulting ROI volumes were 200–203 mm³ for the hippocampus and 100–101 mm³ for the precentral gyrus; the corresponding radii were 3.70–4.12 mm and 3.79–5.73 mm, respectively. The anatomical parcels defined the target locations and clipping boundaries, whereas the smaller parcel-clipped spheres defined the ROIs used for optimization and evaluation.

### 2.5 Montage optimization

Montages for the hippocampal and M1 targets were selected using a two-stage optimization procedure with the MNI152 model provided by SimNIBS [6, 7]. Candidate electrode positions followed the international 10–10 electroencephalography system [16]. Component electric fields for candidate electrode pairs were calculated in SCIRun5 [17]. All possible combinations of two electrode pairs and current amplitudes ranging from 0 to 2 mA were searched in MATLAB.

For each ROI, Pareto optimization was used to find the montage that achieved a mean TIS field of at least 0.2 V/m in the ROI while minimizing the volume of brain tissue outside the ROI exceeding the same field level [4]. The threshold of 0.2 V/m was informed by neurophysiological measurements in cortical network preparations, which established this magnitude as the lower physical limit required to modulate membrane potentials and entrain neuronal activity [18, 19]. This value was used as a numerical optimization target and was not treated as a validated biological threshold for human TIS. Table 3 reports the selected electrode pairs and current amplitudes for each target.

**Table 3:** Optimized stimulation parameters. Electrode configurations and current amplitudes used for targeting the left hippocampus and right primary motor cortex (M1). The montages were selected via exhaustive search optimisation using the MNI152 head model. All electrodes were modeled as cylinde<u>rs with</u> 20 mm diameter and 2 mm height.

| Target | Pair | Electrode locations | Current (mA) |
| --- | --- | --- | --- |
| Left hippocampus | 1 | F8–P8 | 2.00 |
| Left hippocampus | 2 | T7–P7 | 1.59 |
| Right M1 | 1 | Fp2–F6 | 2.00 |
| Right M1 | 2 | C4–CP2 | 0.63 |

**Table 4:** Electrical conductivity values. Isotropic conductivity values were assigned to each tissue and material type in the FEM models. The same values were used consistently across all participants, stimulation targets, and simulation conditions.

| Tissue or material | Conductivity (S/m) |
| --- | --- |
| White matter | 0.126 |
| Grey matter | 0.276 |
| Cerebrospinal fluid | 1.650 |
| Compact bone | 0.008 |
| Spongy bone | 0.025 |
| Scalp | 0.465 |
| Eye tissue | 0.500 |
| Blood | 0.600 |
| Muscle | 0.160 |
| Saline | 1.400 |

### 2.6 TIS field calculation

Each run started from one base head mesh imported into SimNIBS. For pair 1, two cylindrical electrodes (20 mm diameter, 2 mm thickness) were placed on the mesh surface, and the electric field **E**_1_ was calculated using the locations and current amplitudes listed in Table 3. This process was repeated for pair 2 to obtain **E**_2_. The two vector fields were then combined to compute the maximum TIS envelope amplitude [1]. For each element, **E**_1_ and **E**_2_ were ordered such that ||**E**_1_|| *≥* **E**_2_||, and their relative sign was chosen so that the angle *α* between them was acute. The maximum envelope amplitude, *E*_TI_, was calculated as:

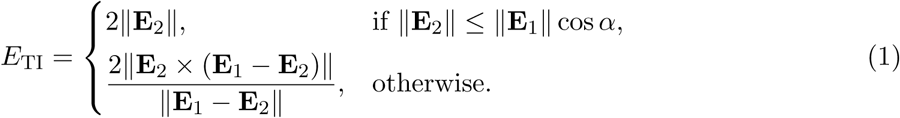

The elementwise *E*_TI_values were interpolated to the participant’s T1-weighted voxel grid and restricted to brain tissue. For each run, 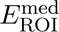 denotes the spatial median of *E*_TI_within the ROI, and 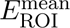 denotes the corresponding spatial mean. 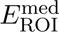 was the primary outcome for the repeatability and ranking analyses. The complementary bootstrap analysis used 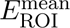 because montage optimization targeted a spatial mean.

Target summaries were calculated from the finite voxels within each spherical ROI. Interpolation and boundary masking left at most 6 of 200–203 hippocampal voxels and 1 of 100–101 M1 voxels without a finite value. Non-finite boundary voxels were excluded elementwise during spatial median and mean calculations; every run retained at least 97% of the hippocampal region and 99% of the M1 region.

### 2.7 Field-repeatability analysis

TIS-field repeatability was quantified separately for each participant, target, and condition. The 40 run-level values of 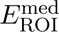 were summarised by their arithmetic mean, sample standard deviation (SD), and coefficient of variation (CV). We calculated CV as 100 *×* SD*/*mean and defined the relative reduction in SD achieved by fixing the base head mesh as 100 *×* (1 *−* SD_fixed_*/*SD_remesh_). Together, these metrics describe both the absolute spread of predicted fields across runs and its magnitude relative to each participant’s mean target field.

### 2.8 Single-run ranking analysis

The ranking analysis was designed to estimate the rank uncertainty associated with any single simulation run. Participants were ordered by their reference mean 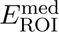 across all 40 runs. For a given pair of participants, the empirical all-pairs ordering-reversal proportion was calculated as the fraction of all 40 *×* 40 = 1,600 observed run combinations in which the participant with the lower reference mean equalled or exceeded the participant with the higher reference mean. Ties were counted as reversals. This proportion describes how frequently a one-run comparison drawn from the observed run distributions disagrees with the ordering based on the complete 40-run means.

Whole-cohort rank stability was assessed using 20,000 Monte Carlo draws. In each draw, one remesh run was selected independently for every participant. Kendall’s tau compared the resulting rank order with the 40-run reference across all 45 participant pairs:

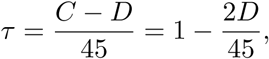

where *C* is the number of concordant pairs and *D* is the number of reversals [20]. A score of +1 indicates that every pair retained its reference order, 0 denotes equal proportions of concordant and reversed pairs, and *−*1 indicates complete rank inversion. This descriptive measure summarises overall cohort stability, whereas the pair-specific proportions identify specific comparisons that are difficult to distinguish reliably. Neither quantity represents a formal hypothesis test.

### 2.9 Bootstrap analysis across run counts

A bootstrap analysis examined how averaging additional remesh runs changed the estimated sampling precision of 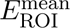. The analysis included all 10 participants and both targets, with 40 run-level values available for each participant–target combination. For each candidate run count from 2 to 40, 20,000 bootstrap samples were drawn with replacement from the 40 observed run values. The arithmetic mean of each bootstrap sample was evaluated against the 40-run reference mean for that participant and target.

Two relative uncertainty measures summarised each bootstrap distribution. The first was half the width of its central 95% interval, expressed as a percentage of the 40-run mean. The second was the 95th percentile of the absolute difference between each bootstrap sample mean and the 40-run mean, again divided by that reference mean and expressed as a percentage. Both measures were retained separately for each participant. Because sampling was performed with replacement, a 40-run bootstrap sample did not necessarily contain each observed run once; uncertainty therefore remained above zero at a sample size of 40. These curves quantify sampling precision relative to the empirical 40-run estimate rather than measuring numerical error relative to an exact continuum solution.

### 2.10 Mesh and tissue-composition outcomes

Element-count analysis evaluated whether total mesh size varied across independent remesh runs. The total tetrahedral element count was extracted from every generated base head mesh. To highlight within-participant variation, element counts were centred on the mean of each participant’s 40-run distribution.

Tissue-composition analysis examined changes in discretised compartment volumes across runs. For each run, the volumes of all tetrahedral elements sharing a given tissue label were summed. The corresponding tissue-specific mean across the 40 remesh runs was subtracted from each run-specific volume, yielding absolute volume deviations in mm^3^. This absolute representation avoids the compositional constraints and artificial anti-correlations inherent to percentage tissue fractions. For each target, the representative participant shown in figures was selected post hoc based on having the largest sum of tissue-specific standard deviations in a preliminary volume-fraction screening. Because this selection used a different metric from the absolute deviations plotted, the displayed cases serve as illustrative examples rather than formal group medians or maximal cases.

## 3 Results

### 3.1 Target-field repeatability across runs

Figure 1 shows 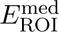 per run across all 10 participants for both targets and numerical conditions. In the remesh condition, hippocampal CVs ranged from 1.81% to 3.65% (median, 2.40%), and M1 CVs ranged from 1.62% to 2.79% (median, 2.54%). Across participants, the observed within-participant range of 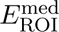 across the 40 remesh runs was 0.0169–0.0322 V/m for the hippocampus and 0.0120–0.0266 V/m for M1 (Table S1).

**Figure 1:**
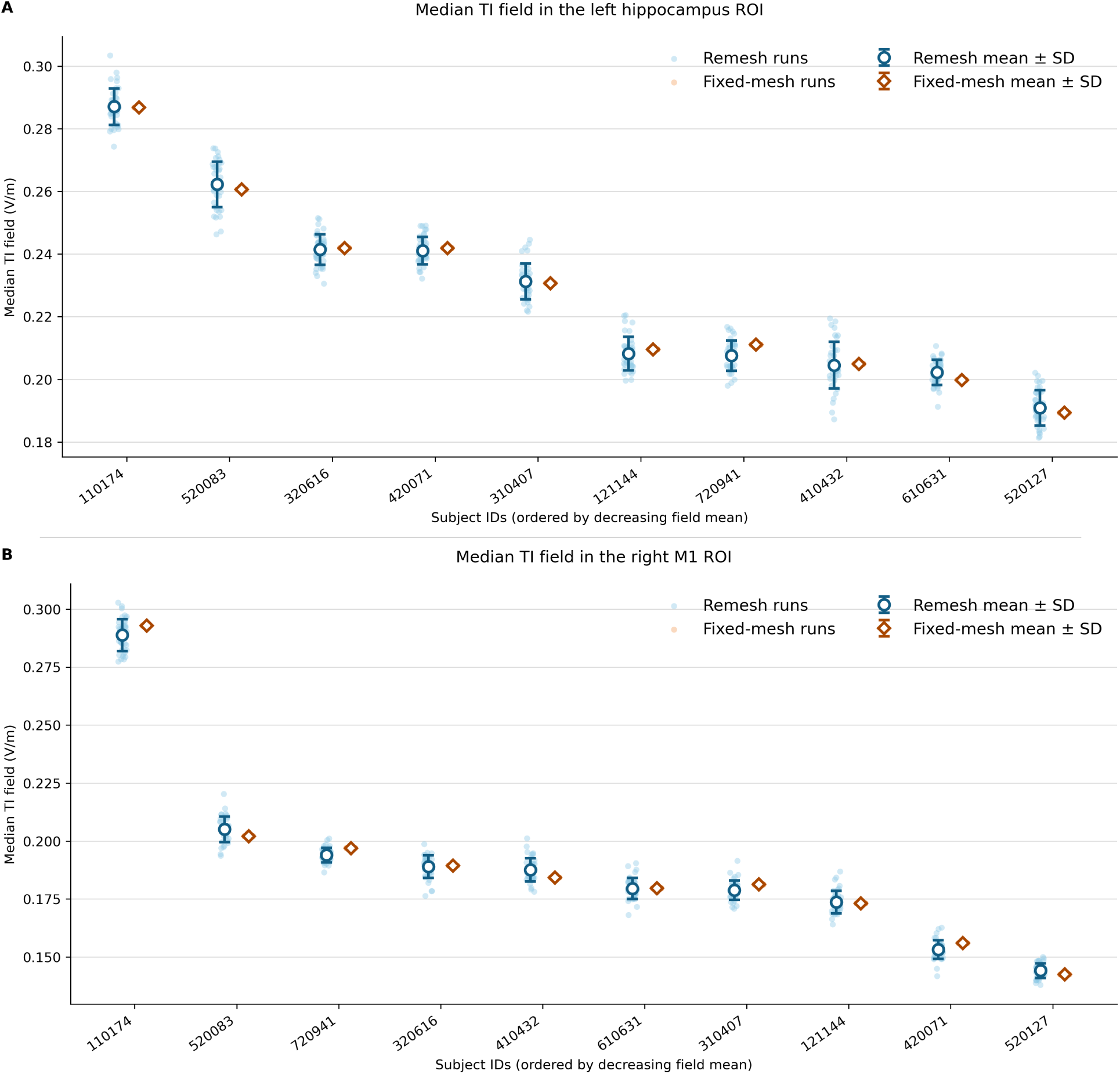
Target-field repeatability across runs. Variation of the median TIS field in the ROI, 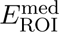, across 40 runs for (A) the left hippocampus and (B) right M1. Pale dots represent individual runs. Open circles (remesh condition) and open diamonds (fixed-mesh condition) show condition means, with error bars denoting *±*1 SD. Individual run points for the fixed-mesh condition are tightly clustered beneath the diamond mean markers. Participants are ordered along the horizontal axis by descending 40-run reference mean 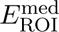.

On the selected fixed meshes, downstream numerical variation was virtually eliminated, yielding a maximum CV of 0.0066% for the hippocampus and 0.0225% for M1. Relative to the corresponding remesh SD, fixing the base head mesh reduced the run-to-run SD by at least 99.6% for every hippocampal model and at least 99.1% for every M1 model. These descriptive ratios are conditional on the selected geometry and do not represent percentages of total numerical variance explained. When run-level distributions of 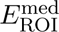 were evaluated against the 0.2 V/m reference threshold, individual runs fell on both sides of this boundary for five hippocampal participants and three M1 participants. For these individuals, applying a threshold-based classification to a single simulation run yielded opposing binary outcomes depending solely on the stochastic mesh realization.

### 3.2 Single-run participant-ordering uncertainty

Figure 2 evaluates cohort rank stability when participant ordering is derived from a single remesh run rather than the 40-run reference mean of spatial medians (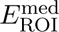) Across 20,000 Monte Carlo selections of one remesh run per participant, median Kendall’s tau relative to the 40-run reference mean ordering was 0.867 (interquartile range, 0.822–0.911) for the left hippocampus and 0.911 (interquartile range, 0.867–0.956) for right M1 (Figure 2B; Table S2). At least one pairwise rank inversion occurred in 95.7% of hippocampal draws and 88.5% of M1 draws, averaging 2.76 and 2.04 pair reversals per draw across the 45 pairwise comparisons, respectively.

**Figure 2:**
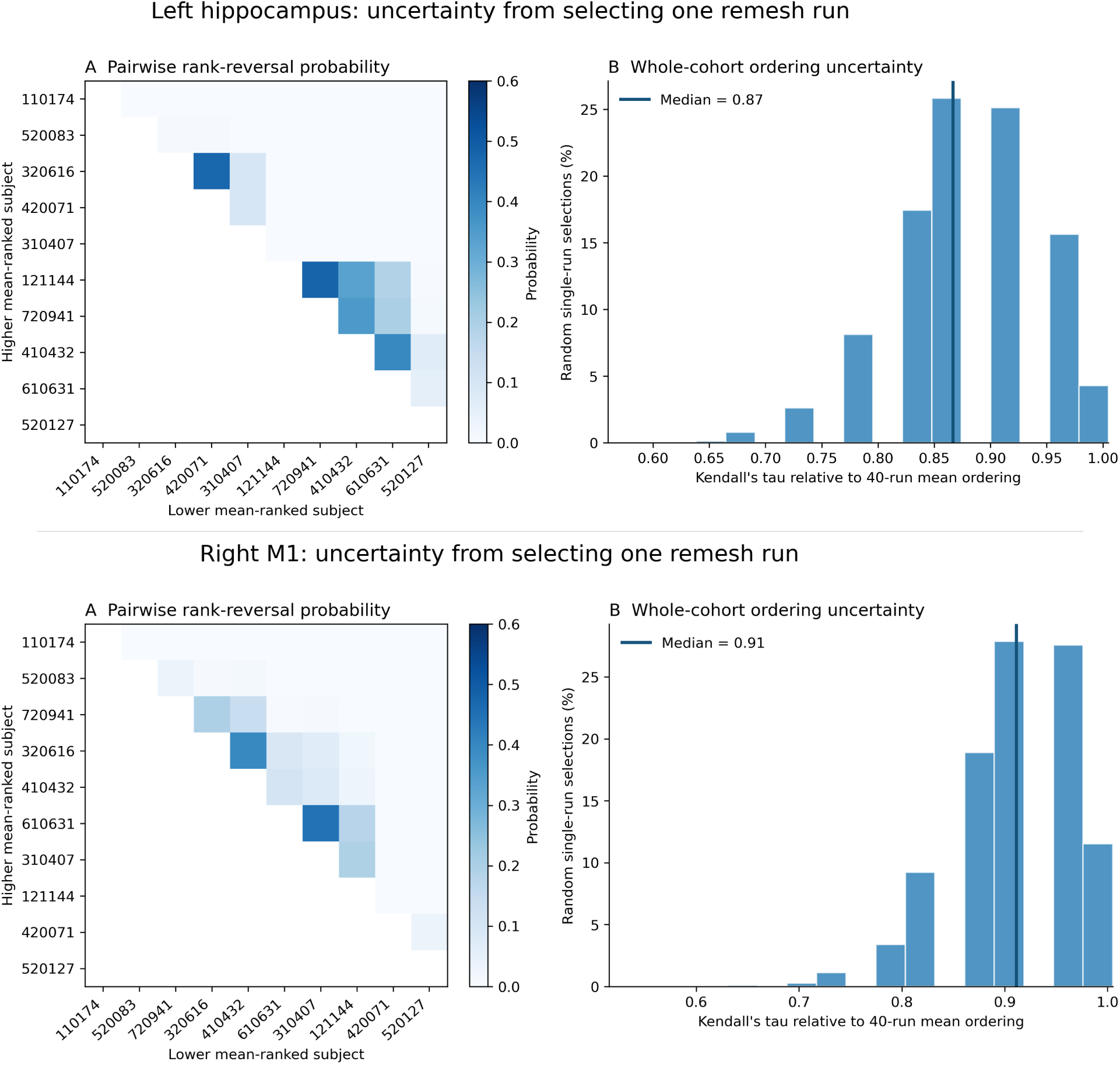
Single-run participant-ordering uncertainty. Evaluation of cohort rank stability for (top row) the left hippocampus and (bottom row) right M1 when drawing a single remesh run per participant. All ranking analyses are based on the run-level spatial median field strength (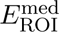). Within each row, panel A shows the empirical pairwise rank-reversal probability matrix across all 45 participant pairs relative to the 40-run reference mean ordering (ordered from highest to lowest 40-run mean 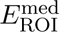). Panel B depicts whole-cohort ordering uncertainty, shown as the distribution of Kendall’s tau across 20,000 random single-run selections relative to that reference ordering. The vertical solid line indicates the median Kendall’s tau value.

The empirical pairwise heatmaps (Figure 2A) demonstrate that rank-reversal probabilities were strictly localized along the sub-diagonal. Reversals were confined to adjacent or closely ranked individuals whose 40-run reference means differed by small margins relative to their meshing variability, while reversal probabilities for distant participant pairs were 0.0%. The largest hippocampal reversal proportion reached 47.9% (for participants whose reference means differed by 0.00060 V/m), while the maximum M1 reversal proportion reached 44.7% (for a reference mean difference of 0.00079 V/m; Table S3).

### 3.3 Bootstrap precision across run counts

Figure 3 shows how sampling uncertainty in the average spatial mean field, 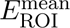, diminished as additional remeshed runs were included. Each participant’s 40-run arithmetic mean served as the empirical reference value. For two-run averages (*n* = 2), the central 95% interval half-width ranged from 2.49–4.80% across hippocampal models and 2.24–3.90% across M1 models. The corresponding 95th percentiles of absolute relative difference ranged from 2.52–4.67% and 2.20–3.94%, respectively.

**Figure 3:**
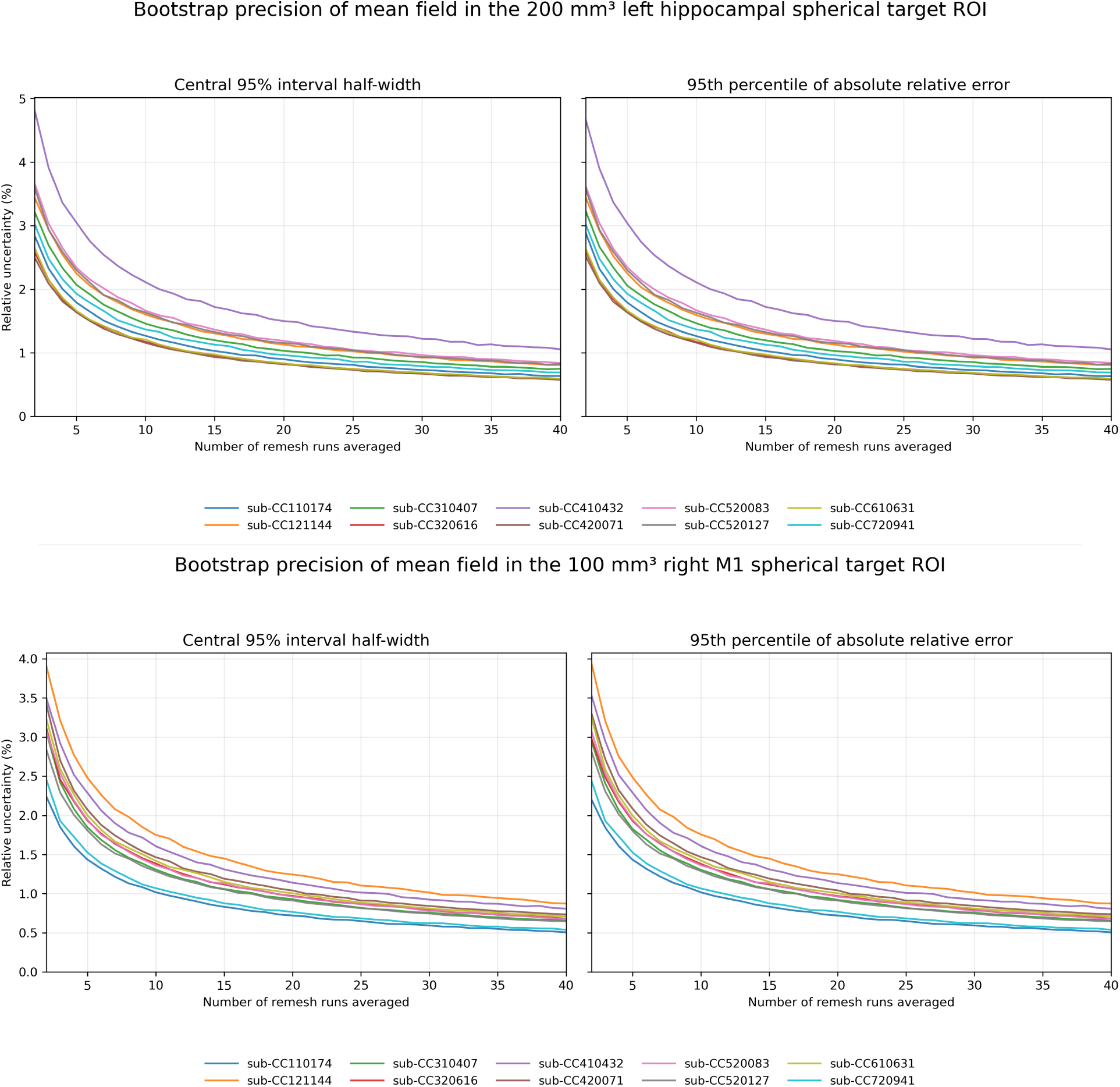
Bootstrap precision across run counts. Sampling precision of the spatial mean TIS field, 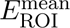, evaluated for (top row) the left hippocampus and (bottom row) right M1, as a function of the number of averaged remesh runs. At each run count (n = 2 to 40), 20,000 bootstrap samples were drawn with replacement from each participant’s 40 remesh runs and averaged. Within each row, the left panel shows half the central 95% bootstrap interval width relative to the 40-run reference mean, and the right panel shows the 95th percentile of absolute relative difference from that empirical reference mean.

Every participant-specific curve followed a characteristic 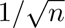 decay as sample size increased. At a bootstrap sample size of *n* = 40, central 95% interval half-widths narrowed to 0.57–1.06% for the hippocampus and 0.51–0.87% for M1. Absolute relative differences yielded nearly identical ranges of 0.58–1.05% and 0.51–0.87%. Values remained strictly above zero at *n* = 40 because resampling with replacement can draw specific runs multiple times while omitting others. This downward trajectory was consistent across all participants and both target regions.

### 3.4 Tetrahedral element-count variability

Figure 4 displays the run-to-run variation in total tetrahedral element count, centred on each participant’s 40-run mean element count. Average mesh size ranged from 2.54 to 3.49 million elements across participants. Across 40 remesh runs, the total within-participant element-count range was 21,344–45,697 elements for the hippocampal experiment and 25,357–48,788 elements for the M1 experiment. Expressed relative to each participant’s mean mesh size, these ranges represented 0.65–1.36% and 0.86–1.40% of total elements, respectively, demonstrating that mesh-size variation was minor relative to total head-model volume.

**Figure 4:**
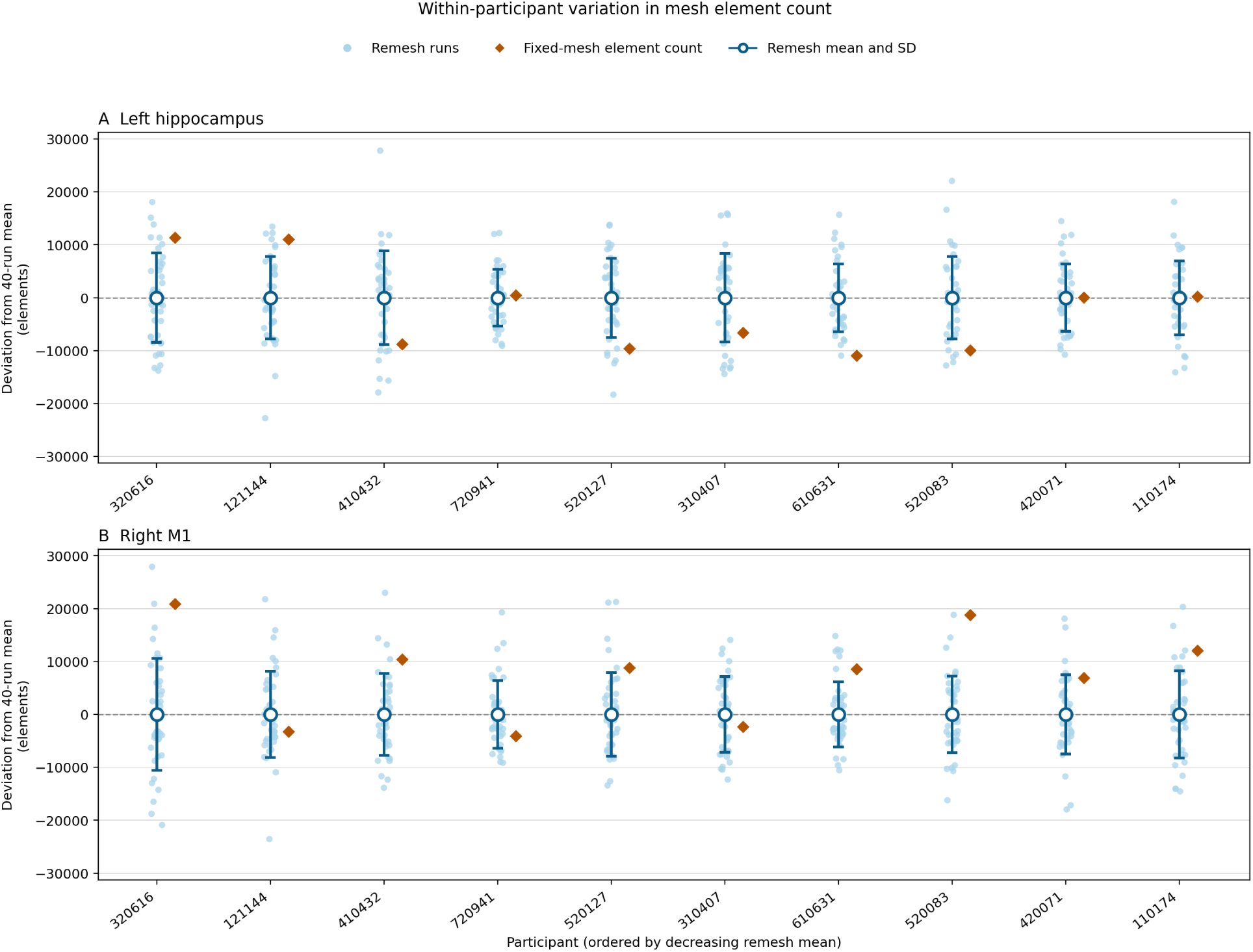
Tetrahedral element-count variability. Within-participant variation in total tetrahedral element count across 40 runs for (A) the left hippocampus and (B) right M1. Individual blue points show deviations from each participant’s 40-run mean element count. Open circles denote the remesh mean (zero on this centred scale), and error bars indicate *±*1 SD. Orange diamonds mark the element-count deviation of the selected fixed mesh. Participants along the horizontal axis are ordered from left to right by decreasing remesh mean.

The element counts of the saved meshes selected for the fixed-mesh condition (orange diamonds in Figure 4) illustrate that selecting a mesh based on field strength does not select for mesh granularity. Although each fixed mesh was chosen because its target field was closest to the 40-run median field, its element count did not systematically coincide with the median or mean element count of the run distribution. Instead, the selected meshes exhibited varying positive and negative deviations from their respective 40-run element-count means.

### 3.5 Tissue-composition variability

Figure 5 shows absolute tissue-volume deviations across 40 remesh runs for the selected high-variability examples. In the hippocampal experiment, participant CC320616 exhibited the largest volume ranges in blood (1,076 mm^3^) and scalp (1,024 mm^3^), followed by grey matter (541 mm^3^), compact bone (521 mm^3^), white matter (496 mm^3^), and CSF (416 mm^3^). Individual runs clearly illustrated local volume swapping between neighbouring tissue compartments; most notably, runs 34 and 40 showed increased scalp volume coinciding with decreased blood volume.

**Figure 5:**
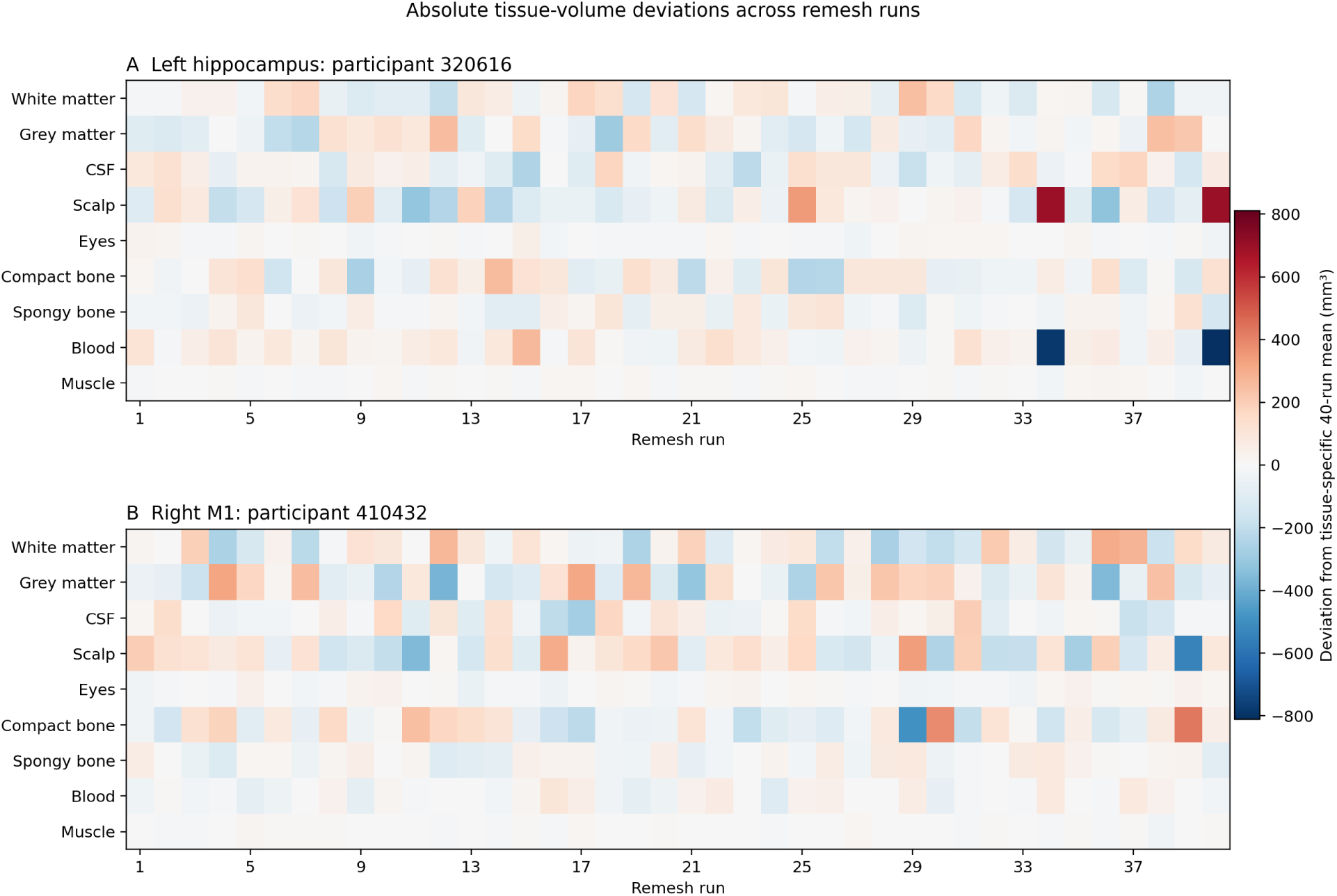
Tissue-composition variability. Absolute tissue-volume deviations across 40 remesh runs for selected illustrative participants from (A) the left hippocampal experiment and (B) the right-M1 experiment. Each cell indicates the difference in mm^3^ between a run-specific compartment volume and its corresponding 40-run mean. Red hues indicate a higher meshed volume than the mean, whereas blue hues indicate a lower meshed volume.

In the M1 experiment, participant CC410432 exhibited the largest volume ranges in compact bone (913 mm^3^) and scalp (883 mm^3^), followed by grey matter (701 mm^3^), white matter (565 mm^3^), and CSF (478 mm^3^). Individual runs again demonstrated compartment swapping; for instance, runs 29 and 39 showed reciprocal shifts between compact bone and scalp volume.

These representative cases demonstrate that stochastic tetrahedralization alters discretised tissue compartment volumes even when generated from an identical tissue-label map. These examples describe specific local variations and do not establish the population-level frequency of specific compartment volume trade-offs or isolate a single tissue as the primary driver of TIS field variability.

## 4 Discussion

This study evaluated the run-to-run repeatability of temporal interference stimulation field predictions when re-executing an identical workflow under stochastic remeshing versus downstream execution on a single fixed mesh. Independent remeshing produced substantially greater repeated-run variation than repeated downstream execution on a selected geometry. Across both target regions, remesh CVs ranged from 1.6% to 3.7%, whereas fixing the base geometry reduced run-to-run standard deviation by more than 99% in every participant, rendering downstream variability virtually zero. This contrast explicitly demonstrates that numerical variability in predicted electric fields is driven almost entirely by the stochastic mesh generation step, rather than by instability in the finite element solver, floating-point differences, or downstream field interpolation. The same qualitative pattern was observed consistently across both deep and superficial target regions.

Differentiating stochastic mesh variability from traditional mesh convergence is essential for interpreting workflow stability. Mesh-convergence and benchmarking studies evaluate how algorithmic choices or spatial refinement alter field distributions relative to a theoretical continuum limit [5, 10, 6]. The present findings demonstrate that even when software pipelines, parameters, and segmentations are held entirely constant, inherent stochasticity in tetrahedral mesh generation introduces an inescapable noise floor. Recognizing this spread as intra-workflow variance—rather than inter-method discrepancy—redefines how numerical precision should be established: pipeline repeatability cannot be assumed solely because the underlying segmentation map and pipeline settings are fixed.

A percent-level coefficient of variation becomes critically important when experimental conclusions depend on small field differences or fixed cutoffs. As demonstrated by the distribution of single-run medians spanning the 0.2 V/m neuromodulation threshold, relying on a single simulation run can alter binary classification decisions regarding whether a participant achieved target engagement. This single-run sensitivity presents a clear vulnerability when ranking closely matched stimulation montages, comparing similar participant cohorts, or conducting prospective target validation, even when no explicit threshold is applied.

Cohort ordering uncertainty was driven primarily by local rank inversions between participants with similar field distributions. While high median Kendall’s tau values indicate that whole-cohort structure is broadly preserved, the high omnibus probability of observing at least one rank reversal (88.5–95.7%) highlights the risk of relying on a single simulation run. Rather than reflecting widespread cohort-wide disruption, this instability is concentrated entirely among adjacent or closely ranked pairs whose reference mean differences are smaller than their numerical meshing variance. Consequently, pair-specific reversal probabilities approaching 50% demonstrate that single-run simulations cannot reliably resolve small individual differences, posing a challenge for studies that rank participants or stratify cohorts based on subtle field variations.

The bootstrap analysis demonstrates that averaging independently meshed simulations effectively mitigates stochastic numerical noise, providing a practical framework for determining multi-run ensemble sizes. The largest precision gains occurred when increasing sample size from 1 to 5 runs, which reduced relative uncertainty bounds from over 4.5% to below 2.0% across all participants and targets. Extending the ensemble average to 10 runs brought the central 95% interval half-width below 1.5% for every model (1.09–1.48% for the hippocampus; 0.88–1.21% for M1). Beyond 10 runs, marginal gains in precision diminished significantly relative to the linear increase in computational execution time. These findings suggest that averaging 5 independent remesh runs provides an efficient baseline for routine applications, whereas averaging 10 runs delivers a highly stable field estimate for studies investigating subtle inter-individual or inter-condition contrasts. Because these precision bounds were evaluated using the spatial mean (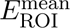) relative to a 40-run empirical reference, they define dataset-specific precision trade-offs rather than a universal requirement. Nevertheless, multi-run averaging offers a clear computational strategy to suppress stochastic meshing variance whenever high-precision field estimates are required.

The tissue-volume analyses confirm that stochastic remeshing alters the underlying spatial discretisation of the head model, providing a potential numerical mechanism for observed field fluctuations. The pipeline used in this study, CHARM, performs direct 3D tetrahedralisation, relying on CGAL algorithms that non-deterministically insert nodes and optimise element shapes across tissue boundaries. Consequently, voxels along tissue transitions are partitioned differently across runs, producing localized volume trade-offs such as the scalp-blood shift in the hippocampal target experiment or the scalp-bone shift in the M1 target experiment. Because tissue conductivities differ substantially across these boundaries, subtle shifts in interface geometry alter local current flow paths. This mechanism highlights an important structural distinction between meshing paradigms. Direct volume-meshing frameworks like CHARM and ROAST are particularly susceptible to boundary-node shifting. By contrast, surface-based meshing pipelines such as SimNIBS headreco construct explicit 2D surface boundaries prior to volumetric tetrahedralisation. Because the interfaces separating distinct conductivity domains are explicitly bounded in surface-based approaches, they may experience substantially less boundary-volume trade-off and lower resulting field variability than direct 3D volume-meshing pipelines. Direct empirical comparisons between direct volume-meshing and surface-constrained paradigms will be required in follow-up work.

### 4.1 Limitations

The numerical estimates in this study come from a deliberately narrow design comprising ten head models, two targets, one segmentation approach, and one simulation pipeline. Other anatomical regions, electrode geometries, tissue maps, or meshing algorithms may produce different levels of variation. The reported percentages should therefore be interpreted as specific measurements of this workflow rather than universal properties of finite element modelling or temporal interference stimulation.

The experiment measured repeatability, not accuracy. No analytical solution or highly refined reference mesh was available, so the study cannot determine the numerical bias of any realisation. Element counts and tissue-volume deviations also do not measure local element quality or demonstrate mesh convergence. High repeatability on a fixed mesh does not guarantee accuracy, nor does variation among valid remesh realisations reveal which estimate lies closest to the underlying continuum solution.

The primary repeatability analysis evaluated spatial medians (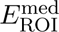) within parcel-clipped target spheres, whereas secondary analyses introduced operational variations in spatial support and metrics. Specifically, fixed-mesh selection was computed across the full anatomical parcel rather than the clipped sphere, and the fixed-mesh condition evaluated downstream execution on only one selected geometry per participant–target combination. Consequently, the fixed-mesh condition estimates downstream repeatability conditional on those specific geometries rather than yielding a complete separation of between-mesh and within-mesh variance.

Similarly, the bootstrap analysis evaluated spatial means (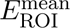) across resampled run counts relative to the observed 40-run mean. While spatial means and full-parcel field distributions correlate tightly with local target medians, these implementation differences mean the resulting precision curves describe the sampling behavior of ensemble-averaged spatial means. They provide a complementary analysis of estimator precision rather than formal confidence intervals for single-run 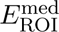 values or a validated stopping rule.

Interpolation left a small number of non-finite voxels along certain ROI boundaries. Every run retained at least 97% of hippocampal support and 99% of M1 support, but spatial summaries were not restricted to a common finite mask across all runs. Minor changes in the included boundary voxels may therefore contribute a small portion of the observed field variation. Finally, the tissue-composition figures illustrate specific high-variability cases to demonstrate possible boundary trade-off mechanisms, but they do not establish the population-level frequency of those specific compartment patterns across the full cohort.

## 5 Conclusions

Independent remeshing introduced percent-level variation in target TIS field estimates across both the left hippocampus and right M1. Repeated downstream execution on a single fixed geometry produced substantially less spread, demonstrating that workflow variability was driven almost entirely by non-deterministic mesh generation rather than solver instability or downstream post-processing. A single independently meshed realisation preserved broad cohort-level differences but frequently inverted the rank order of closely matched participant pairs. Furthermore, a complementary bootstrap analysis showed that averaging multiple independent remesh runs progressively improved the sampling precision of target TIS field estimates, with five to ten runs providing an effective balance between numerical stability and computational cost. These findings quantify single-workflow repeatability rather than absolute numerical accuracy. Stochastic mesh variation should be evaluated, controlled, or mitigated through multi-run averaging whenever experimental conclusions depend on subtle field differences or fixed neuromodulation thresholds.

## Acknowledgements

**BI:** Conceptualization, Methodology, Investigation, Software, Data curation, Formal analysis, Visualization, Writing – original draft, and Project administration. **SR:** Conceptualization, Methodology, Software, Validation, Investigation, Writing – review & editing, Supervision. **MA:** Writing – review & editing, Funding acquisition. **JT:** Writing – review & editing. All authors reviewed and approved the final version of the manuscript.

## Funding

BI was supported by EPSRC iCASE, Perspectum and the University of Sheffield under reference number EP/W524360/1. JT and MA were supported by an Advanced Research and Invention Agency (ARIA) grant to MA with number SCNI-PR01-P08. SR was supported by a National Institute of Neurological Disorders and Stroke grant to SR with number R01NS133229.

## Conflicts of interest

The authors declare no competing interests.

## Author contributions

### Ethics statement

This numerical study used previously acquired CamCAN anatomical data under the dataset’s access and ethics provisions. Data collection and sharing for this project was provided by the Cambridge Centre for Ageing and Neuroscience (CamCAN). CamCAN funding was provided by the UK Biotechnology and Biological Sciences Research Council (grant number BB/H008217/1), together with support from the UK Medical Research Council and University of Cambridge, UK.

## Supplementary Information

**Table S1:** Participant-level repeatability of the median electric-field strength within each region of interest across 40 simulation runs. Results are shown for the left hippocampus (A) and right primary motor cortex (M1; B). For each participant, the table reports the mean and standard deviation (SD) obtained when the mesh was independently regenerated for each simulation (remesh) and when the same mesh was reused across simulations (fixed mesh). The coefficient of variation (CV) describes the relative run-to-run variability in the remeshed simulations, while SD reduction shows how much this variability decreased when a fixed mesh was used.

A. Left hippocampus
| Subject | Remesh (V/m) |  | Fixed mesh (V/m) |  | SD reduction (%) | Remesh CV (%) |
| --- | --- | --- | --- | --- | --- | --- |
|  | Mean | SD | Mean | SD |  |  |
| CC110174 | 0.28704 | 0.00580 | 0.28674 | < 0.000005 | 100.0 | 2.02 |
| CC121144 | 0.20820 | 0.00533 | 0.20954 | < 0.000005 | 100.0 | 2.56 |
| CC310407 | 0.23124 | 0.00573 | 0.23069 | < 0.000005 | 100.0 | 2.48 |
| CC320616 | 0.24144 | 0.00487 | 0.24191 | < 0.000005 | 100.0 | 2.02 |
| CC410432 | 0.20453 | 0.00747 | 0.20494 | < 0.000005 | 100.0 | 3.65 |
| CC420071 | 0.24109 | 0.00437 | 0.24192 | 0.00002 | 99.6 | 1.81 |
| CC520083 | 0.26222 | 0.00724 | 0.26058 | < 0.000005 | 100.0 | 2.76 |
| CC520127 | 0.19092 | 0.00563 | 0.18934 | < 0.000005 | 100.0 | 2.95 |
| CC610631 | 0.20222 | 0.00402 | 0.19981 | < 0.000005 | 100.0 | 1.99 |
| CC720941 | 0.20761 | 0.00480 | 0.21106 | < 0.000005 | 100.0 | 2.31 |

B. Right M1
| Subject | Remesh (V/m) |  | Fixed mesh (V/m) |  | SD reduction (%) | Remesh CV (%) |
| --- | --- | --- | --- | --- | --- | --- |
|  | Mean | SD | Mean | SD |  |  |
| CC110174 | 0.28878 | 0.00689 | 0.29286 | 0.00001 | 99.8 | 2.39 |
| CC121144 | 0.17368 | 0.00485 | 0.17310 | < 0.000005 | 100.0 | 2.79 |
| CC310407 | 0.17877 | 0.00419 | 0.18125 | < 0.000005 | 100.0 | 2.35 |
| CC320616 | 0.18894 | 0.00486 | 0.18940 | < 0.000005 | 100.0 | 2.57 |
| CC410432 | 0.18755 | 0.00502 | 0.18424 | < 0.000005 | 100.0 | 2.68 |
| CC420071 | 0.15319 | 0.00407 | 0.15596 | 0.00004 | 99.1 | 2.66 |
| CC520083 | 0.20504 | 0.00548 | 0.20206 | < 0.000005 | 100.0 | 2.67 |
| CC520127 | 0.14416 | 0.00313 | 0.14244 | < 0.000005 | 99.9 | 2.17 |
| CC610631 | 0.17956 | 0.00452 | 0.17965 | < 0.000005 | 100.0 | 2.52 |
| CC720941 | 0.19395 | 0.00314 | 0.19696 | < 0.000005 | 100.0 | 1.62 |

*Note.* Each row summarizes 40 runs per condition. Mean and sample standard deviation (SD) were calculated from the run-level values of $E_{ROI,med}$ . Values shown as < 0.000005 V/m fall below the reporting precision. CV is $100 \times SD/mean$ . SD reduction is $100 \times (1 - SD_{fixed}/SD_{remesh})$ .

**Table S2:** Uncertainty in subject ordering when one remesh run is selected per subject.

| ROI | Median $\tau$ | $\tau$ Q1 | $\tau$ Q3 | Any reversal (%) | Mean reversals per draw | Maximum pair probability (%) |
| --- | --- | --- | --- | --- | --- | --- |
| Left hippocampus | 0.867 | 0.822 | 0.911 | 95.7 | 2.76 | 47.9 |
| Right M1 | 0.911 | 0.867 | 0.956 | 88.5 | 2.04 | 44.7 |

*Note.* Kendall's $\tau$ compares each of 20,000 random single-run orderings with the ordering based on the 40-run means of $E_{\text{ROI},\text{med}}$ . A value of +1 indicates identical ordering, 0 indicates equal proportions of concordant and reversed subject pairs, and $-1$ indicates complete reversal. Q1 and Q3 are the 25th and 75th percentiles of $\tau$ . Any reversal is the percentage of draws in which at least one of the 45 pairs changed order. Mean reversals per draw is the average number of changed subject pairs among the 45 comparisons. Maximum pair probability is the largest exact reversal probability across all subject pairs. These quantities describe numerical uncertainty and are not p-values.

**Table S3:** Subject pairs with the highest single-run rank-reversal probabilities.

| ROI | Rank | Higher-mean subject | Mean $E_{\text{ROI}}^{\text{med}}$ (V/m) | Lower-mean subject | Mean $E_{\text{ROI}}^{\text{med}}$ (V/m) | Reversal probability (%) |
| --- | --- | --- | --- | --- | --- | --- |
| Left hippocampus | 1 | CC121144 | 0.20820 | CC720941 | 0.20761 | 47.9 |
| Left hippocampus | 2 | CC320616 | 0.24144 | CC420071 | 0.24109 | 47.1 |
| Left hippocampus | 3 | CC410432 | 0.20453 | CC610631 | 0.20222 | 39.6 |
| Left hippocampus | 4 | CC720941 | 0.20761 | CC410432 | 0.20453 | 35.4 |
| Left hippocampus | 5 | CC121144 | 0.20820 | CC410432 | 0.20453 | 33.1 |
| Right M1 | 1 | CC610631 | 0.17956 | CC310407 | 0.17877 | 44.7 |
| Right M1 | 2 | CC320616 | 0.18894 | CC410432 | 0.18755 | 39.4 |
| Right M1 | 3 | CC720941 | 0.19395 | CC320616 | 0.18894 | 19.5 |
| Right M1 | 4 | CC310407 | 0.17877 | CC121144 | 0.17368 | 19.4 |
| Right M1 | 5 | CC610631 | 0.17956 | CC121144 | 0.17368 | 17.6 |

*Note.* Subjects are ordered by the mean $E_{\text{ROI}}^{\text{med}}$ across their 40 remesh runs. For each pair, reversal probability is the fraction of all $40 \times 40 = 1,600$ run combinations in which the lower-mean subject equals or exceeds the higher-mean subject. The five largest probabilities are shown for each target.

## Notes

### Competing Interest Statement

The authors have declared no competing interest.

